# From cytoskeletal dynamics to organ asymmetry: a non-linear, regulative pathway underlies left-right patterning

**DOI:** 10.1101/052191

**Authors:** Gary S. McDowell, Suvithan Rajadurai, Michael Levin

## Abstract

Consistent left-right asymmetry is a fundamental aspect of the bodyplan across phyla, and errors of laterality form an important class of human birth defects. Its molecular underpinning was first discovered as a sequential pathway of left-and right-sided gene expression that controlled positioning of the heart and visceral organs. Recent data have revised this picture in two important ways. First, the physical origin of chirality has been identified; cytoskeletal dynamics underlie the asymmetry of single cell behavior and of patterning of the left-right axis. Second, the pathway is not linear: early disruptions that alter the normal sidedness of upstream asymmetric genes do not necessarily induce defects in the laterality of the downstream genes or in organ *situs*. Thus, the LR pathway is a unique example of two fascinating aspects of biology: the interplay of physics and genetics in establishing large-scale anatomy, and regulative (shape-homeostatic) pathways that correct errors of patterning over time. Here, we review aspects of asymmetry from its intracellular, cytoplasmic origins to the recently-uncovered ability of the LR control circuitry to achieve correct gene expression and morphology despite reversals of key “determinant” genes. We provide novel functional data, in *Xenopus laevis*, on conserved elements of the cytoskeleton that drive asymmetry, and repair of downstream gene expression anomalies over developmental time. LR patterning can thus serve as a paradigm of how subcellular physics and gene expression cooperate to achieve developmental robustness of a body axis.

## 1 Introduction

Most vertebrates (and many invertebrates) have bilaterally-symmetric external bodyplans. Yet these same animals exhibit consistent asymmetries in the position or anatomy of internal organs such as heart, viscera, and brain [1]. Defects in LR asymmetry are an important class of human birth defects, including heterotaxia (the lack of concordance between internal organs, allowing each organ to individually decide on its placement on the left or right side of the body), single organ inversions such as dextrocardia (the reversal in position and morphology of the heart), and isomerisms (symmetry of the LR axis, leading to either duplication or complete loss of unpaired organs such as the spleen). Patients with complete reversal of asymmetry (*situs inversus*) have fewer health consequences than these other conditions because heterotaxia and isomerisms often involve inappropriate connections between the heart, lungs and other visceral organs [2, 3, 4, 5, 6]. Interestingly, consistent laterality affects not only the asymmetric organs, but also the manifestation of numerous diseases and conditions affecting paired (seemingly symmetrical) organs, such as hip dysplasia, limb defects, and eye development [7, 8, 9, 10, 11]. This includes location and incidence of tumors [12, 13, 14, 15, 16, 17], immune responses [18, 19], and innervation of sensory organs such as eyes [20].

LR asymmetry is a unique puzzle in development [21], in complex organisms and even in colonies of simpler ones [22]. The embryonic anterior-posterior (AP) axis probably exists in order to place sense organs at the end of the animal that encounters novel environments first (oriented with the main axis of motion). The dorso-ventral (DV) axis can be set by gravity or sperm entry point. However, once those two orthogonal axes are set, the alignment of the LR axis is fixed; in order to distinguish L from R, symmetry has to be broken. Crucially, it is not merely a question of making L different from R, but doing it so that the LR axis is consistently oriented with respect to the AP and DV axes. The former process results in fluctuating asymmetry (an indicator of stress via the difficulty of keeping the two halves of the body precisely coordinated during growth); the latter is true biased asymmetry of anatomical structures. This is a difficult problem, as our universe does not macroscopically distinguish left from right [23, 24]. This problem was noted long ago, by workers studying chiral biochemistry and its implications for development [25, 26, 27], clinical observations of mirror-imaging in anatomical features of human twins [28], unidirectional coiling of snail shells [29], and functional handedness in neural lateralization [30]. Since then, consistent LR asymmetry has been identified not only in anatomical structures from somites [31] to hair whorls [32, 33, 34, 35, 36] to deer antlers [37], but also in functional features such as immune response [38, 39, 40] and cancer [41, 42, 17, 43]. Thus, the mechanisms of LR patterning are not only of fundamental interest in developmental biology, but are central to many interesting questions of physiology and evolution.

Early mechanistic work in this field identified a set of chemical agents that was able to perturb (randomize) asymmetry in animal models [44, 45, 46, 47, 48]. The first molecular explanations for the asymmetry of body organs came from studies in the chick [49], with the identification of asymmetrically-expressed genes, such as the left-sided *Sonic hedgehog* (*Shh*) and *Nodal*, the inductive and repressive relationships among these genes, and functional studies showing that aberrant expression of any of these was sufficient to randomize the *situs* of the heart, gut, and other viscera [50, 51]. These data not only helped explain organ laterality in normal development but also provided a mechanistic explanation for laterality disturbances long known to occur in conjoined twins [52]. The central component of the LR pathway was the left-sided cassette formed by *Shh* inducing expression of *Nodal* inducing expression of *Lefty* inducing expression of *Pitx2* [53, 54, 55].

The conserved molecular pathway in *Xenopus*, chick, mouse, and zebrafish as described in early work on LR asymmetry [56, 57] has since been extended considerably, as described in Figure 1 and reviewed in [58, 59], by loss-and gain-of-function approaches in a range of species that reveal the functional connections among LR patterning proteins. New players include Activin, Follistatin, Derrire, Coco, Mad3, BMP, Noggin4, and Fgf8 [60, 61, 62, 63, 64, 65, 53, 66, 67, 68, 69, 70]. Recently, other downstream factors have been increasingly demonstrated in unilateral function during LR determination, such as calcium signaling and retinoic acid [71, 72, 73, 74, 75, 76, 77, 78, 79]. Together, this body of work reveals a progressive cascade of left-and right-specific activities that involve powerful signaling pathways. These signaling molecules then provide distinct signals to organ primordia on either side of the midline, resulting in asymmetric organogenesis.

**Figure 1:**
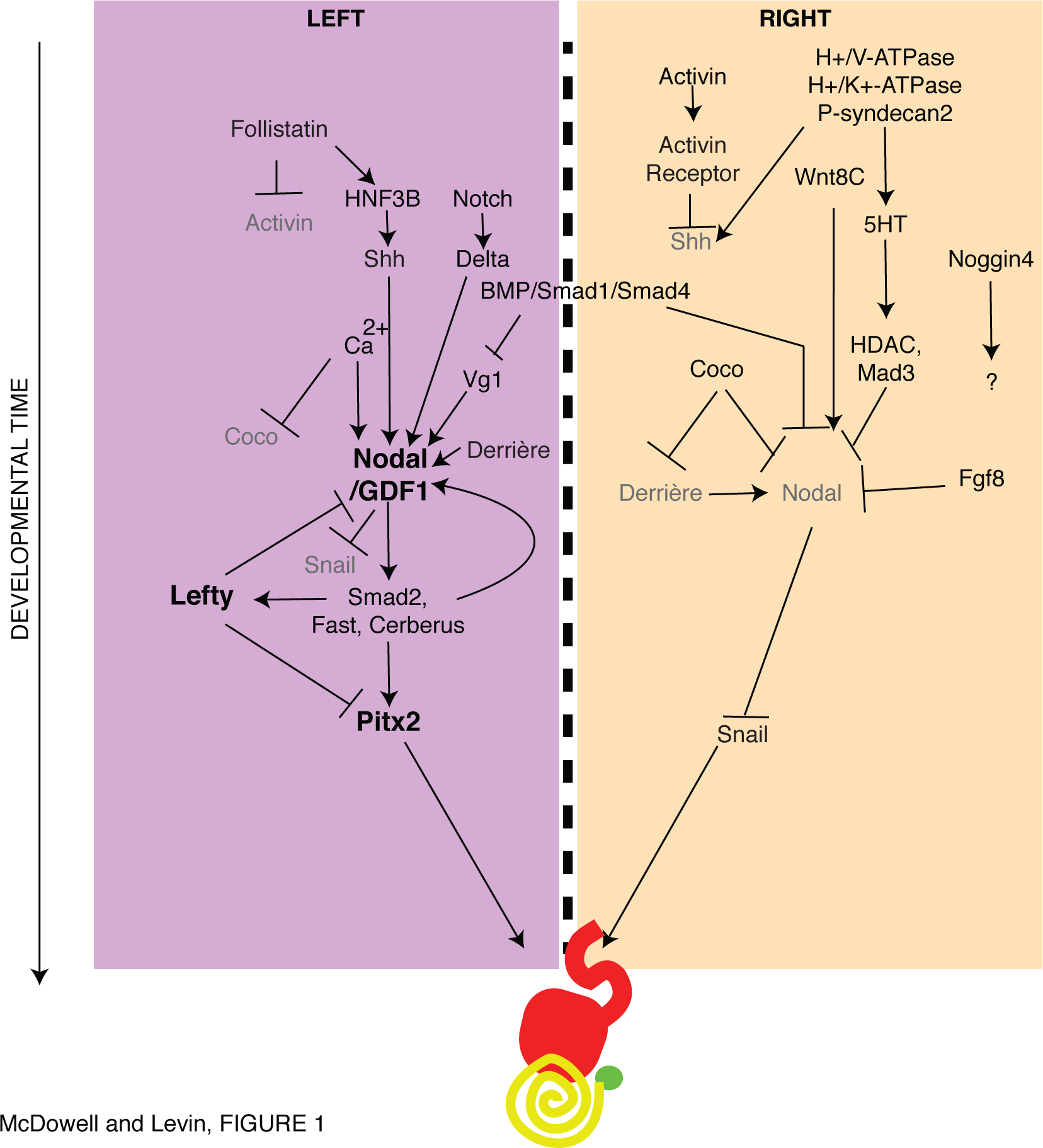
Classical view of progressive linear cascades in the asymmetry-determining pathway. Numerous molecular components have now been implicated in the establishment of L or R identity of the two embryonic halves. The functional data testing knockdown and over-expression of each gene, followed by examination of the expression of downstream steps, have given rise to a view of the pathway as a linear progressive cascade of repressions and inductions, with variations between species (for example, the activation of *Nodal* by Notch/Delta represents the system in mouse and chick, but Notch/Delta actually inhibits *Nodal* in zebrafish and sea urchin). A core cassette is the promotion of left by left-sided *Sonic hedgehog* (*Shh*) expression inducing *Nodal* and then *Lefty* and *Pitx2*, which ultimately sets the *situs* of the heart, while Activin repression of *Shh* on the right of the node promotes right with the inactivation of *Nodal* and therefore derepression of *Snail*. Such a linear perspective predicts that defects in the expression of upstream genes will result in corresponding defects in their downstream targets, propagating errors towards organ heterotaxias. Recent data have indicated that this view of asymmetry is limited in 2 ways. Gene regulatory networks on their own can not distinguish spatial properties (cannot tell L from R or constrain geometry in any way), nor can they directly control the physical forces needed to shape organ morphogenesis. Moreover, a linear pathway does not correctly predict the many examples of progressive defect repair in the LR signaling that leads to organ *situs*. Thus, any such transcriptional network must be bookended by chiral physical elements upstream (to link the very first asymmetric transcriptional event to one side but not the other) and downstream (to shape the asymmetric organs); it must also include mechanisms for sensing aberrant gene expression and correcting downstream steps.

### 1.1 Physics upstream and downstream of transcriptional networks

Regardless of the completeness of a transcriptional pathway or network model, one has to explain why the first asymmetric gene becomes expressed on one side but not the other. It is a fundamental limitation of transcriptional regulatory networks that they do not in themselves constrain geometry — no gene regulatory network (GRN) can generate consistently chiral output (distinguish left from right). Thus, upstream of any such regulatory cascade must lie some piece of physics which is able to break symmetry and reliably orient subsequent events with respect to the other two axes [80, 81].

Interestingly, this is not strictly a multicellular phenomenon, in a number of systems, from bacteria to ciliates to human somatic cells in vitro, single cells exhibit consistent chirality in their movements and behaviors [25, 82, 83, 84, 85, 86, 87, 88, 89, 90, 91, 92, 93, 94, 95], both as individual cells and as collectives [96, 92]. For example, outgrowth of neurites is predominantly clockwise [84, 85, 97, 90]. The asymmetric growth and migration of various cell types on micropatterned surfaces has demonstrated the role of actomyosin networks in the generation and maintenance of consistent chirality of migration, cell shape, and tissue morphogenesis [82, 91]. Thus, metazoan organisms face the problem of amplifying subcellular chirality into tissue-wide asymmetry across a midline, but do not need to reinvent the symmetry-breaking and orientation steps, as these appear to be ancient and ubiquitous.

One candidate for a chiral element of physics upstream of asymmetric gene expression is ciliary flow at neurulation [72, 98, 99]; this model however faces numerous problems which have been detailed elsewhere [100, 101, 102, 103, 104]. Recent functional data revealed conserved mechanisms in a range of organisms from plants to mammals, that establish asymmetry without a ciliated structure, or long before it forms; indeed, many phyla including some vertebrates determine their LR axis very early after fertilization [105, 106, 107, 108, 109, 110, 111, 112, 113, 114, 115]. Even mouse embryos are known to exhibit molecular and functional asymmetries (e.g., components such as cofilin, which is also asymmetric in cleavage-stage frog embryos) as early as the cleavage stages [107, 116, 117].

While ciliary flow may impinge on downstream transcriptional events in those species where it exists (not pig [108] or chick [118, 119] for example), it cannot be the origin of asymmetry in most phyla. From data in a range of model species, it is clear that numerous aspects of development, including maternal protein localization [120, 121, 111, 122], Wnt signaling [123], and small signaling molecules [114] are already consistently asymmetric long before cilia appear: most animal embryos can tell their Left from their Right at very early stages. Thus, the search for the *origin* of asymmetry has been extended far upstream of neurulation [124, 47].

One interesting class of models links asymmetry to chromatid segregation [125, 126, 127, 128] a proposal that needs to be tested in the available model systems, as it offers the possibility of linking asymmetry to the fundamental dynamics of DNA. This model has been linked to data on birth defects and epithelial morphogenesis in humans and mice [129]. As in *Xenopus*, the mouse model has also revealed that α-tubulin organization is critical for asymmetry, via studies of the protein Mahogunin [130, 105, 111, 112, 131].

A remarkably prescient prediction was made by a paper published before all of the molecular work on the LR pathway [132], in which Brown and Wolpert hypothesized a chiral element that initiates biased transport inside the early embryonic blastomeres. More recent work in several models have confirmed early hypotheses while elaborating on these ideas [133, 134], showing that the cytoskeleton is centrally involved in generating the original asymmetric cues that break and orient symmetry. A number of model species such as *C. elegans* [135, 74, 136, 137, 138] and snails [139, 140, 141, 142] were known to use cytoskeletal dynamics to determine chiral cell behavior and subsequent LR patterning. However recent data in these models, together with mammalian cells and *Drosophila* [143, 144], have revealed many of the key details.

The intriguing structure of centrioles [145], including their anti-clockwise rotation [146], and the role of microtubules in generating asymmetry in neutrophils [147, 93], plants [148], and frog [111] and *C. elegans* embryos [149, 150], lends evidence to the theory that the centrosome could be a symmetry-breaking, chiral structure [151, 100]. Evidence for intrinsic chirality, and not interactions with the substrate, were provided by the counter-clockwise rotation exhibited in zebrafish melanophores [94]. A particularly illustrative case of spontaneous intracellular chiral cytoskeletal organization again illustrates the role of actomyosin networks in the ability of α-actinin to spontaneously arrange directionally, or reverse the directionality if the protein is grossly overexpressed [89].

Studies in *Drosophila* have demonstrated the effect of intrinsic cellular chirality on embryonic laterality [143, 152, 153, 154, 155, 156, 157, 158, 159, 160]. Unconventional myosins, such as Myosin1d, have a clear role in affecting the asymmetry of the gut and genitalia in *Drosophila* [158, 159] and this asymmetry is due to a direct effect on the actin filaments in epithelia [154]. Actin motility on Myo1c occurs in counterclockwise direction [161]; myosin V is a left-handed spiral motor toward the plus end of actin [162], while myosin II is right-handed spiral motor [163]. Moreover, the effect of these unconventional myosins on organismal asymmetry is linked to their effect on intrinsic cellular chirality [160], and individual cells can contribute to mechanical differences in generating chirality at the tissue level [153, 160], a finding also demonstrated in *Caenorhabditis elegans* [113, 134].

Our own work in *Xenopus* has demonstrated a number of key features that point to the importance of the cytoskeleton at multiple points in laterality. Firstly, fundamental cytoskeletal proteins such as tubulins and myosins are functionally important for normal embryonic laterality at the very earliest points of embryonic development, immediately post-fertilization [111, 112]. The cytoskeleton appears to be required for the normal early localization of asymmetric components such as ion channels [121, 111, 122], a mechanism widely conserved to many somatic cell types [164, 165]. Secondly, the conservation of phenotypes obtained from mutations in cytoskeletal and cytoskeleton-associated proteins exhibited in frog, compared to those of plants [111, 112], snails [105], and *Drosophila* [112], demonstrate the need for physical signals upstream of gene regulatory networks (GRNs): for example, the organ-specific effects of myosins on *Drosophila* laterality were replicated in frog [112], transcending the differences in molecular LR pathways between vertebrates and invertebrates and raising the question of how the cytoskeleton generates chirality. Finally, the apparent disconnect between the asymmetric expression of *Nodal* and subsequent organ *situs* observed in mouse [130] and replicated in frog [112] highlight a conserved role for the cytoskeleton not only in laterality, but in the ability to correct earlier defects in laterality between the point of expression of markers of laterality and the positioning of visceral organs.

Downstream of the chiral physics within cells lie physiological mechanisms that amplify subcellular chiralities into true LR asymmetries across cell fields. One such system is the diffusion-based LALI (local activation, long range inhibition) system described in mouse [166, 167]. Another is the chiral bioelectric gradient that redistributes intracellular morphogens such as maternal serotonin [120, 168, 169, 170, 171, 110, 172, 78, 173]. This process is most well-understood in frog embryos, but has also been observed and functionally implicated in amphioxus [174], sea urchin [168, 175, 176], *C. elegans* [177, 178, 179], zebrafish [120, 180, 181], and recently humans [182, 183]. Indeed very recent analysis of the differences between blastomeres in the very earliest stages of embryo development, identified distinct metabolites in L vs. R cells in the 8-cell embryo using mass spectrometry [114]. The majority of these metabolites themselves have roles in functional regulation of ion transport [184, 185, 186, 187, 188, 189, 190, 191, 192], suggesting possible feedback loops in electrophysiology that could be important amplifying mechanisms for initially subtle LR asymmetry. Both of these systems impinge upon a key asymmetric gene NODAL [62, 193], and lie upstream of a cascade of asymmetric gene interactions. However, much as it needs a physical process to anchor consistent asymmetry upstream, genetic pathways likewise need to control physical forces in order to actually implement asymmetric morphogenesis.

GRNs feed into specific proteins that harness physical forces such as tension and adhesion to control asymmetric bending and growth of internal organs [194]. Mechanical forces are critical for the rotation and looping of internal organs ([195, 196], see reviews for the heart in chick [197, 198, 199, 200]). In the development of *Xenopus* gut morphology, a rightward torsion results in concave and convex topologies for cells on the left and right sides of the lateral plate mesoderm respectively, and cells on the right elongate twice as much as cells on the left, although proliferation rates remain the same [201]. Similarly, in *Drosophila*, asymmetries arising from planar cell polarity in gut looping through the activity of myosin1d were able to generate differences in tension, with greater tension on the left side of the cell driving leftward rotation [160]. In heart looping, a similar role for actin and myosin in driving dextral looping has been exhibited in zebrafish [196] and chick [202]. Recently computational modeling has supported evidence suggesting that differential growth supplies the forces that cause the heart tube to bend ventrally, while cytoskeletal movements can drive rightward torsion [203]. The maintenance of symmetry could also be due to differences in the composition of the extracellular matrix (ECM) as observed in perturbations of ECM composition in frog and chick embryos [204, 48].

### 1.2 Developmental regeneration: robustness of the LR pathway

Bracketed upstream and downstream by interaction with physical forces, the middle of the LR patterning process consists of a pathway made up of sequentially interacting left-and right-sided gene products (Fig. 1). The implication of this kind of induction/repression model is that if something goes wrong upstream — if a particular gene is expressed on the wrong side for example — then the downstream elements will likewise be incorrect. For example, in the early work on LR determining genes in chick, it was shown that if the left-sided *Shh* gene was misexpressed on the right side, right-sided expression of the normally left-sided gene *Nodal* followed, and organs were subsequently randomized.

Importantly however, development in many species is highly regulative: early errors (e.g., splitting embryos in half [205]) can become subsequently corrected. While many perturbations overcome this basic shape homeostasis property (resulting in birth defects), it is nevertheless true that embryos are highly robust because of the ability to remodel after some kinds of deviations from the normal sequence of events. One example is the derivation of largely normal frog faces from early embryos with severely malformed facial structures, which self-correct over time [206]. The mechanisms of robustness and shape homeostasis are poorly understood at the mechanistic level, although they are now increasing objects of interest in molecular developmental biology [207, 208, 209] and regenerative medicine [210, 211, 212].

Self-correction capabilities in the LR pathway are a novel, molecularly-tractable example of regenerative repair. It is a uniquely-accessible context in which to study the regulatory mechanisms that recognize and reverse abnormal patterning states. Here, we identify new aspects of the cytoskeletal machinery that lie upstream of the LR asymmetry patterning pathway and investigate their robustness. We quantitatively analyze published data as well as our new functional studies to reveal remarkable flexibility in the GRN that belies a simple linear pathway to reveal remarkable pattern robustness.

## 2 Results: endogenous repairing of induced left-right defects

The canonical description of the left-right patterning pathway in vertebrates, regardless of the proposed point of symmetry breaking, follows the linear track of leftward activation of *Nodal*, downstream activation of *Lefty* and *Pitx2*, and thence to the correct placement of visceral organs. However, we have observed in our recent studies of the determination of laterality in *Xenopus*, many anomalies and exceptions to this neat pipeline.

For example, in studying proteins with roles in early embryonic asymmetries [213], and in particular cytoskeletal proteins with a conserved role in determining embryonic laterality [105, 112], the number of embryos with reversed organs is actually smaller than the number with incorrect *Nodal* expression (Table 1). This suggests that the pathway from *Nodal* to organ *situs* is not as linear as had been assumed: having incorrect *Nodal* expression does not actually mean that an embryos organs will be reversed, despite *Nodals* established role as a determinant or master regulator of LR *situs*.

**Table 1:** Effects of protein overexpression on Nodal laterality and organ situs and the degree of repair of incorrect laterality, calculated as the percentage of embryos with incorrect Nodal expression that have correct organ situs, using separate clutches for analysis. R: right; Bi: bilateral; N: none (correct sidedness is Left). A positive degree of repair indicates that embryos with incorrect gene expression went on to have normal organ situs, indicating that some defects in expression of early laterality markers are being corrected by the time of organ placement.

| **Protein** | **Incorrect**<br>***Nodal* expression** | **Reversed Organs** | **Reference** | **Degree of repair** |
| --- | --- | --- | --- | --- |
| α-tubulin T56I | **37%**<br>(R: 10%, Bi: 9%, N: 18%) | **21%** | [112] | **43%** |
| Lis-N99 | **26%**<br>(R: 9%, Bi: 8%, N: 9%) | **18%** | [112] | **30%** |
| Lis-C137 | **21%**<br>(R: 7%, Bi: 9%, N: 5%) | **10%** | [112] | **52%** |
| 14-3-3E | **65%**<br>(R: 12%, Bi: 30%, N: 23%) | **30%** | [112] | **54%** |

The difference between the number of embryos with incorrect *Nodal* expression and reversed organs has been particularly noticeable in our studies on the role of the intracellular motor protein family myosins in laterality, with up to 22-fold differences between the number of embryos misexpressing *Nodal* and the number of tadpoles with incorrect placement of internal organs (Table 2). This suggests again that although a high percentage of embryos exhibit incorrect *Nodal* expression, this does not lead to the reversal of organs that a linear pathway from *Nodal* to organ morphogenesis would suggest. The most obvious example is Myosin1cA, which results in almost half of the embryos having incorrect *Nodal* expression, but the resulting populations organ *situs* is almost normal (only 2% heterotaxia) — a very significant repair capability downstream of Myosin function.

**Table 2:**
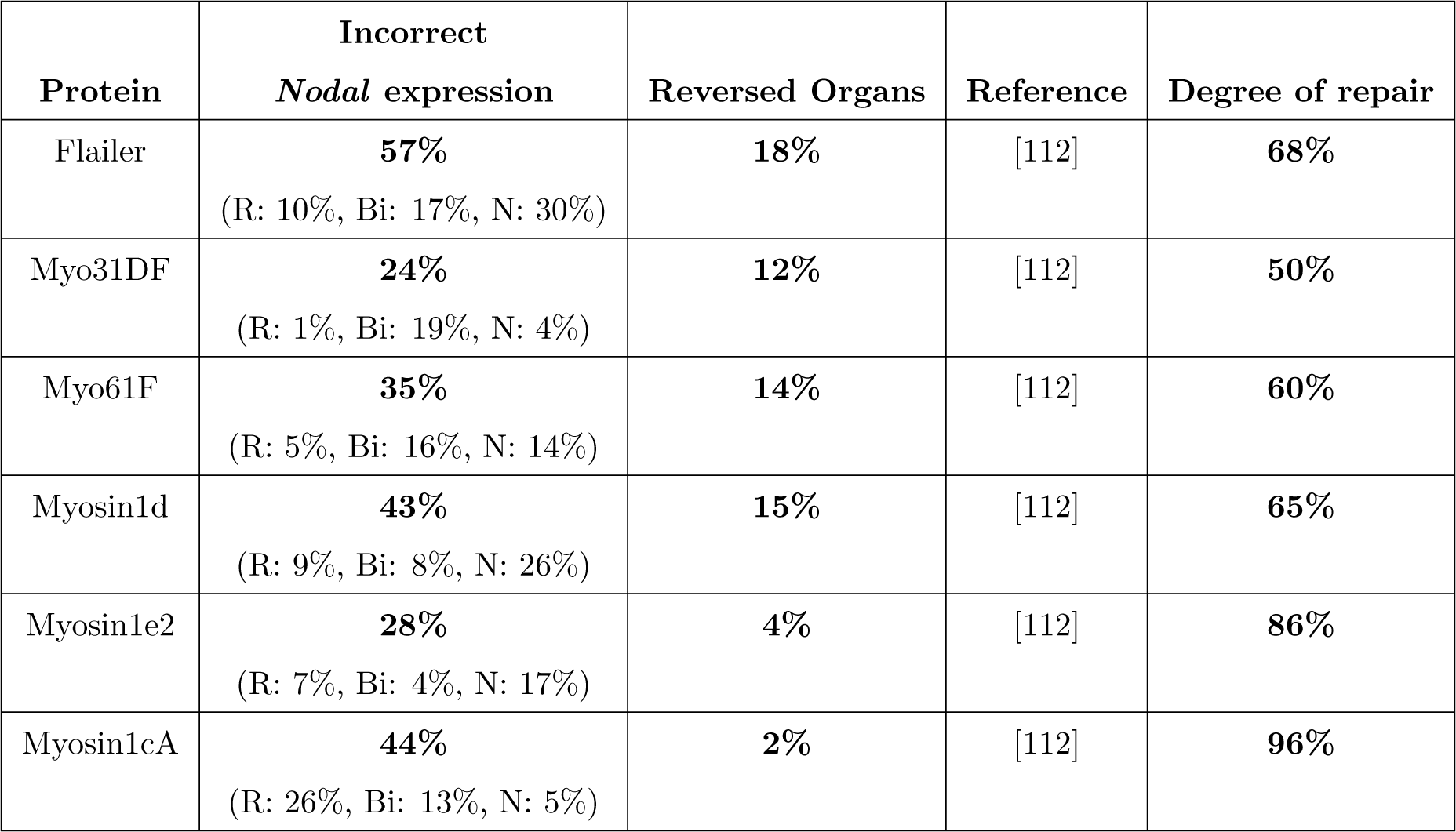
Effects of protein overexpression on Nodal laterality and organ situs and the degree of repair of incorrect laterality, calculated as the percentage of embryos with incorrect Nodal expression that have correct organ situs, using separate clutches for analysis. R: right; Bi: bilateral; N: none (correct sidedness is Left). A positive degree of repair indicates that embryos with incorrect gene expression went on to have normal organ situs, indicating that some defects in expression of early laterality markers are being corrected by the time of organ placement.

This has prompted us to review the literature covering previous experiments in *Xenopus* which cover the effects on both *Nodal* laterality and organ *situs*. Looking at untreated embryos alone has been informative: for example, we find that the level of incorrect *Nodal* expression in untreated embryos, while low, is higher than the level of organ *situs* ([112], Table 3). However, others have found that the elevated level of misplaced *Nodal* matches to a spontaneous rate of cardiac reversal alone of 3-8% [214], which in our hands is an extremely high rate of spontaneous organ reversal for all organs [105, 112], let alone the heart. However, this may be a simple biological difference between frog populations or experimental procedures in laboratories. Regardless, there are still many more embryos observed with *Nodal* discrepancies than organ *situs* misplacement in both cases.

**Table 3:** State of Nodal laterality and organ situs and the degree of repair of incorrect laterality, calculated as the percentage of embryos with incorrect Nodal expression that have correct organ situsin control (untreated) embryos. A positive degree of repair indicates that embryos with incorrect gene expression went on to have normal organ situs, indicating that some defects in expression of early laterality markers are being corrected by the time of organ placement.

| Incorrect *Nodal* expression | Reversed Organs | Reference | Degree of repair |
| --- | --- | --- | --- |
| 12% | 3-8%<br>spontaneous cardiac reversals | [214] | 33-75% |
| 34% | <1% | [186] | 97% |
| 7% | 1% | [112] | 86% |
|  | 0.5% | [105] | N/A |

It is from these discrepancies that we began to formulate our hypothesis of fixing of left-right mispatterning during embryogenesis. Throughout the formation of the left-right axis, we propose that there are numerous checks and balances on the mechanism of generating and maintaining asymmetry. As there are numerous health problems associated with a single organ being misplaced with respect to the orientation of all others, it is incredibly important that all organs be oriented in their correct position with respect to one another. But if all organs are reversed, in the case of *situs inversus totalis*, there are few health problems associated, if any (and it is suggested that there is an under-diagnosis of this condition due to the lack of reported corresponding health concerns [215]). What we observe in nature is not a 50:50 split in organs all on one side or the other, however; across vertebrates, all individuals within a species, and all species, have the same orientation. If organs were not able to position independently of one another, but were all dependent together on the position of *Nodal, Lefty* and *Pitx2* expression, then potentially we would observe a 50:50 split throughout nature as it would not matter which side the set of organs chose to occupy. The fact that this is not the case, and that there is a preferred orientation, suggests that individual organs have the potential to orient independently of one another. Therefore, to prevent organs from doing this, we propose that there is a strong evolutionary pressure to stick with a preferred orientation for organs, and to reinforce that orientation throughout development, as it is crucial to the survival of the organism. If this were the case, we might find that it is difficult to push the level of heterotaxia under a variety of perturbations to high levels, or to flip entirely. This is in fact the case in many scenarios. It would also explain the apparent lack of conservation in mechanisms for left-right patterning across closely-related vertebrates, as different mechanisms could become prominent in different species depending on the nature of their embryonic development. What could be conserved, however, is the importance of the chirality of the cytoskeleton, now a widely-described phenomenon from cells in culture to embryos, and it is the mechanism of how the cytoskeleton can generate, maintain and rescue chirality and chiral defects that would then be of most interest to the left-right field.

Reviewing the literature for evidence of these phenomena, we found some interesting supportive observations. To begin with, there were a number of observations showing various degrees of correction or fixing of laterality defects from early to late markers (Table 4).

**Table 4:** Effects of connexin protein overexpression or drug treatment on Nodal laterality and organ situs and the degree of repair of incorrect laterality, calculated as the percentage of embryos with incorrect Nodal expression that have correct organ situs, using separate clutches for analysis. H7: gap junction communication disruptor. R: right; Bi: bilateral; N: none (correct sidedness is Left). A positive degree of repair indicates that embryos with incorrect gene expression went on to have normal organ situs, indicating that some defects in expression of early laterality markers are being corrected by the time of organ placement.

| **Protein** | **Incorrect**<br>***Nodal* expression** | **Reversed Organs** | **Reference** | **Degree of repair** |
| --- | --- | --- | --- | --- |
| Cx26 2 of 4-cell:<br>ventral | **48%**<br>(R: 1%, Bi: 1%, N: 46%) | **32%** | [186] | **33%** |
| Cx43 2 of 4-cell:<br>ventral | **32%**<br>(R: 0%, Bi: 5%, N: 27%) | **22%** | [186] | **31%** |
| H7 2 of 4-cell:<br>ventral | **57%**<br>(R: 5%, Bi: 14%, N: 38%) | **29%** | [186] | **49%** |

The same was to be found in a variety of drug treatments of early *Xenopus* embryos affecting laterality (Table 5).

**Table 5:**
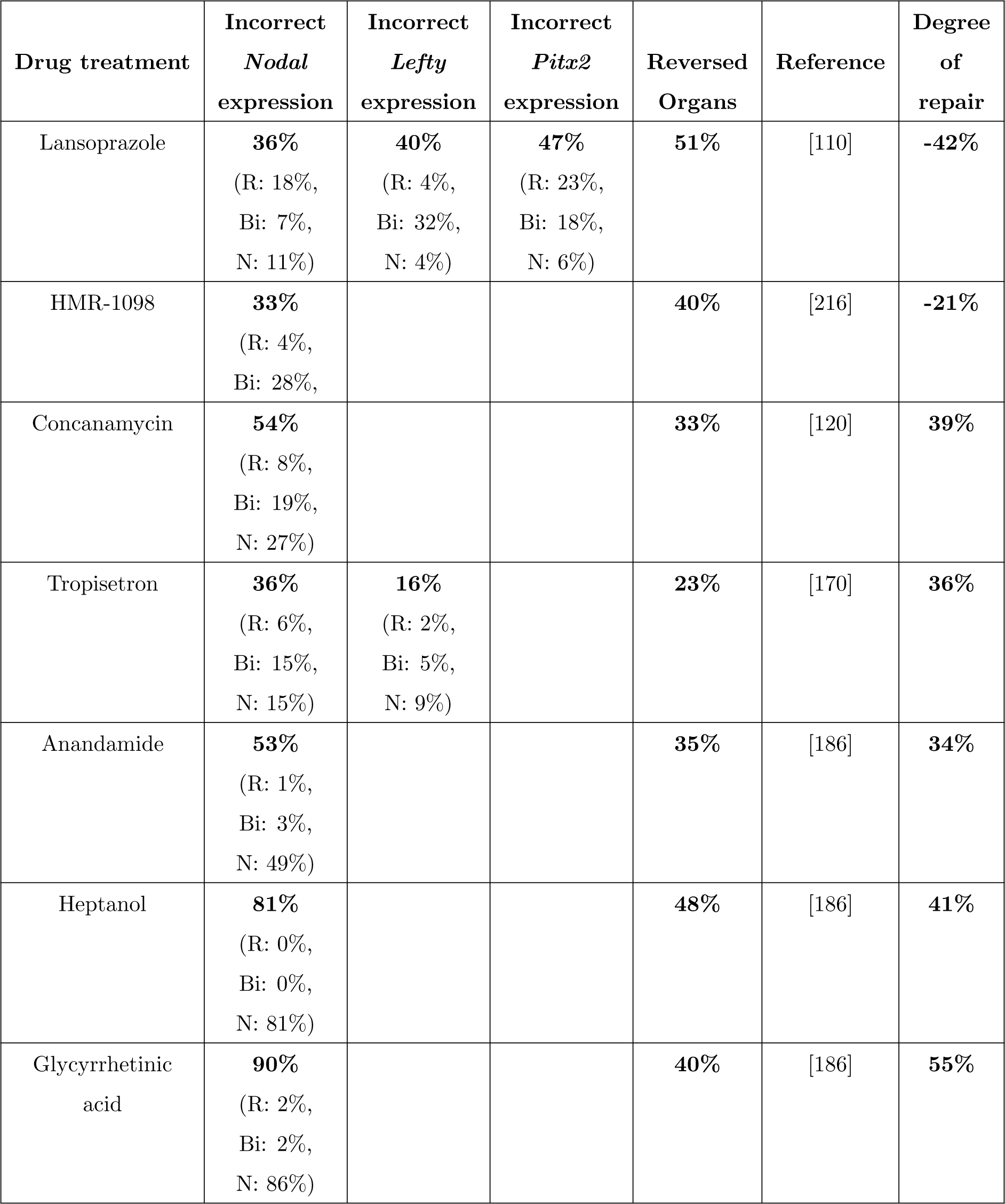
Effects of drug treatment on Nodal laterality and organ situs and the degree of repair of incorrect laterality, calculated as the percentage of embryos with incorrect Nodal expression that have correct organ situs, using separate clutches for analysis. Lansoprazole: blocker of H+/K+-ATPase pump; HMR-1098: blocker of KATP channel; Concanamycin: blocker of H+-V-ATPase pump; Tropisetron: serotonin receptor blocker; Anandamide: gap junction communication blocker; Heptanol: gap junction communication blocker; Glycyrrhetinic acid: gap junction communication blocker. R: right; Bi: bilateral; N: none (correct sidedness is Left). A positive degree of repair indicates that embryos with incorrect gene expression went on to have normal organ situs, indicating that some defects in expression of early laterality markers are being corrected by the time of organ placement. A negative degree of repair indicates that early errors are amplified by subsequent steps.

We found evidence from other research groups (see Table 6) of cases where there is no difference, such as with the injection of BVg1 mRNA into the R3 cell of a 16-cell embryo [217]; in other cases, such as injection of BMP2 mRNA into the R2 cell of the 16-cell embryo caused a small perturbation in *Nodal* laterality but no effect on organ *situs* [217]. In some cases, such as the injection of Xwnt-8 mRNA into the L4 cell of the 16-cell embryo, errors either accumulate past *Nodal* expression or bypass *Nodal* altogether [217]. This is possibly also the case for injection of BMP2 mRNA into the L2 cell of the 16-cell embryo: Hyatt and Yost [217] describe a truncated left phenotype for the expression of *Nodal*, but *Nodal* expression still appears to be correct scoring on laterality alone, and yet the rate of organ reversal is much higher.

**Table 6:** Effects of mRNA and protein injection on Nodal laterality and organ situs and the degree of repair of incorrect laterality, calculated as the percentage of embryos with incorrect Nodal expression that have correct organ situs, using separate clutches for analysis. Degree of repair for Xwnt-8 indicates that defects in early laterality markers are being amplified by the point of organ placement. Degree of repair for BMP2, BVg1 R4 and Activin A indicates that defects in early laterality markers are being corrected by the point of organ placement.

| **Protein** | **Incorrect**<br>***Nodal* expression** | **Reversed Organs** | **Reference** | **Degree of repair** |
| --- | --- | --- | --- | --- |
| BVg1 R3 16-cell | **88%** | **89%** | [217] | **-1%** |
| tAR L1 16-cell | “complex” | **51%** | [217] | **Unknown** |
| BMP2 L2 16-cell | **30%** | **85/58%** | [217] | **Unclear** |
| BMP2 R2 16-cell | **17.5%** | **0%** | [217] | **Complete**<br>**correction** |
| Xwnt-8 L4 | **17%** | **63%** | [217] | **-271%** |
| Xwnt-8 L4, BVg1 R1 | **66%** | **88%** | [217] | **-33%** |
| Xwnt-8 L4, BVg L1 | “not determined” | **14%** | [217] | **Unknown** |
| BVg1 L4 16-cell | **15%** | **10%** | [218] | **33%** |
| BVg1 R4 16-cell | **75%** | **52%**<br>cardiac reversal | [218] | **31%** |
| Activin A PROTEIN<br>Right-sided Injection<br>at Stage 20 | **83%** | **49%** | [219] | **41%** |
| Activin A PROTEIN<br>Left-sided Injection<br>at Stage 20 | Not shown | **19%** | [219] | **Unknown** |

Likewise, there are instances of drug treatments which result in incorrect *Nodal* laterality and organ *situs* but apparently have no effect on the positioning of *Lefty* in the canonical left-right pathway and so affect certain parts of the pathway while leaving others unaffected, such as the examples in Table 7.

**Table 7:** Effects of drug treatment on Nodal and Lefty laterality and organ situs and the degree of repair of incorrect laterality, calculated as the percentage of embryos with incorrect Nodal expression that have correct organ situs, using separate clutches for analysis. GR113808: serotonin receptor blocker; Iproniazid: monoamine oxidase inhibitor. R: right; Bi: bilateral; N: none (correct sidedness is Left). Degree of repair for GR113808 indicates that defects in Nodal and organ situs are the same but that Lefty is bypassed. Degree of repair for Iproniazid indicates that defects in early laterality markers are being corrected by the point of organ placement.

| Drug treatment | Incorrect<br>*Nodal*<br>expression | Incorrect<br>*Lefty*<br>expression | Reversed<br>Organs | Reference | Degree<br>of<br>repair |
| --- | --- | --- | --- | --- | --- |
| GR113808 | **43%**<br>(R: 10%,<br>Bi: 26%,<br>N: 7%) | **14%**<br>(R: 3%,<br>Bi: 2%,<br>N: 9%) | **44%** | [170] | **-2%** |
| Iproniazid | **43%**<br>(R: 10%,<br>Bi: 21%,<br>N: 13%) | **15%**<br>(R: 2%,<br>Bi: 4%,<br>N: 10%) | **30%** | [170] | **30%** |

To test this hypothesis further, we wished to find out whether the differences in levels we observed were due to the comparison of *Nodal, Lefty* and *Pitx2* laterality to organ *situs* from different clutches of embryos. We have data from a selection of cytoskeletal proteins generated by this method in Table 8 from our previous analysis [112].

**Table 8:** Effects of cytoskeletal protein overexpression on Nodal, Lefty and Pitx2 laterality and organ situs and the degree of repair of incorrect laterality, calculated as the percentage of embryos with incorrect Nodal expression that have correct organ situs, using separate clutches for analysis. R: right; Bi: bilateral; N: none. Degree of repair for Mgrn C314D indicates that defects in laterality are accumulating by the point of organ placement. Degree of repair for Mgrn wt and Mgrn G2A indicate that defects in early laterality markers are being corrected by the point of organ placement.

| Drug treatment | Incorrect<br>*Nodal*<br>expression | Incorrect<br>*Lefty*<br>expression | Incorrect<br>*Pitx2*<br>expression | Reversed<br>Organs | Degree<br>of<br>repair |
| --- | --- | --- | --- | --- | --- |
| Control | **7%**<br>(R: 1%,<br>Bi: 2%,<br>N: 4%) | **3%**<br>(R: 0%,<br>Bi: 1%,<br>N: 2%) | **4%**<br>(R: 0%,<br>Bi: 2%,<br>N: 2%) | **1%** | **86%** |
| Mgrn wt | **44%**<br>(R: 10%,<br>Bi: 23%,<br>N: 11%) | **8%**<br>(R: 0%,<br>Bi: 0%,<br>N: 8%) | **4%**<br>(R: 0%,<br>Bi: 0%,<br>N: 4%) | **6%** | **91%** |
| Mgrn G2A | **26%**<br>(R: 4%,<br>Bi: 10%,<br>N: 12%) | **20%**<br>(R: 3%,<br>Bi: 1%,<br>N: 16%) | **23%**<br>(R: 6%,<br>Bi: 10%,<br>N: 7%) | **15%** | **42%** |
| Mgrn C314D | **14%**<br>(R: 1%,<br>Bi: 8%,<br>N: 5%) | **8%**<br>(R: 4%,<br>Bi: 4%,<br>N: 0%) | **14%**<br>(R: 0%,<br>Bi: 7%,<br>N: 7%) | **17%** | **-21%** |
| Ect2-trunc | **24%**<br>(R: 6%,<br>Bi: 8%,<br>N: 10%) | **27%**<br>(R: 10%,<br>Bi: 2%,<br>N: 15%) | **34%**<br>(R: 12%,<br>Bi: 12%,<br>N: 10%) | **25%** | **-4%** |

We therefore used individual clutches and at different timepoints took embryos for *in situ* hybridization against *Nodal, Lefty*, and *Pitx2* and scored the remaining tadpoles for organ *situs*, as we have not observed whether individual clutches of embryos have been analyzed for the within-clutch variation, or whether the variation observed previously is simply clutch-to-clutch variation. As observed in Table 8, the data correspond very well with our previous findings ([112], Table 9).

**Table 9:**
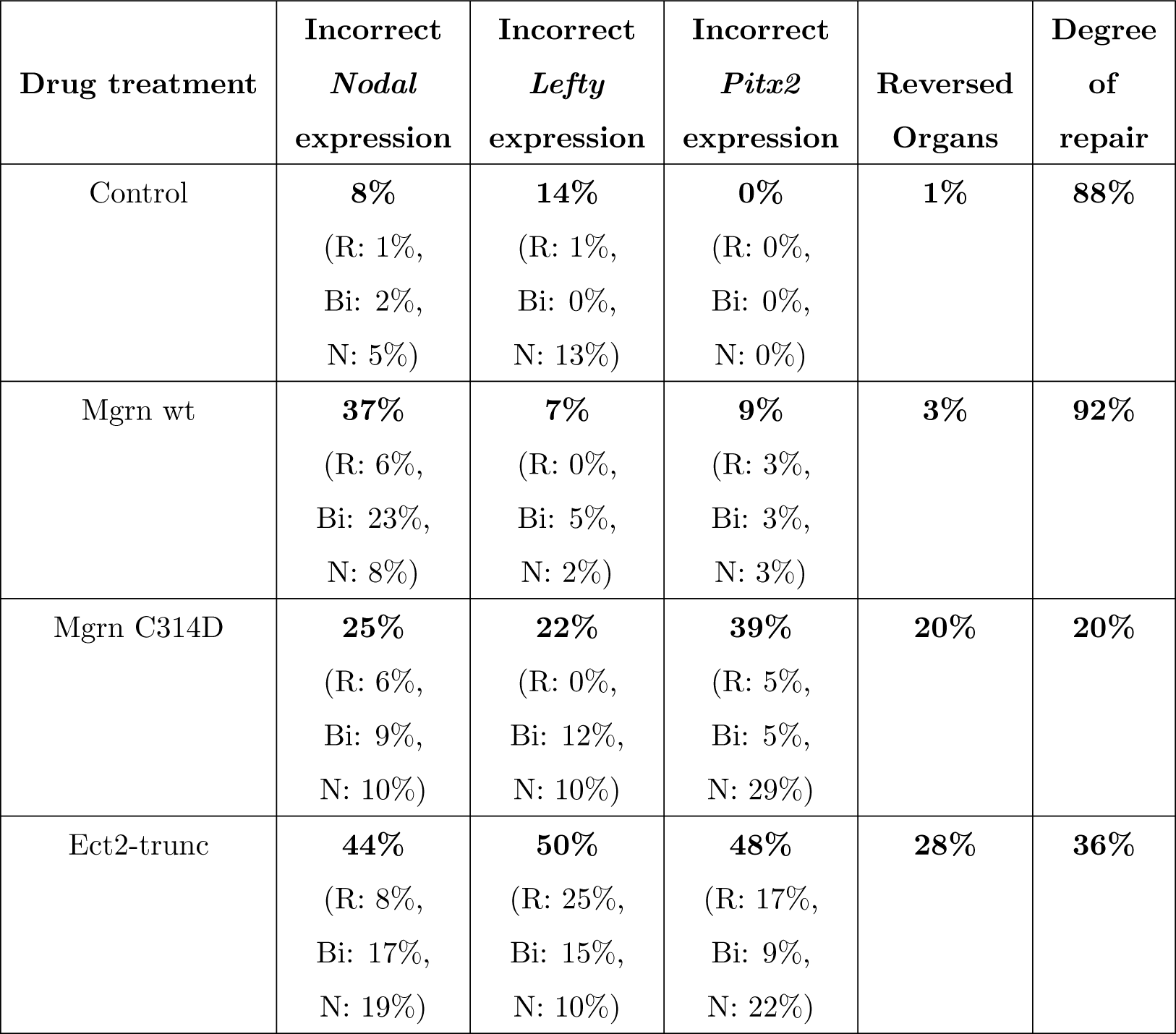
Effects of cytoskeletal protein overexpression on Nodal, Lefty and Pitx2 laterality and organ situs and the degree of repair of incorrect laterality, calculated as the percentage of embryos with incorrect Nodal expression that have correct organ situswithin the same clutches. R: right; Bi: bilateral; N: none. Degree of repair for Mgrn C314D indicates that defects in laterality are accumulating by the point of organ placement. Degree of repair for Mgrn wt indicates that defects in early laterality markers are being corrected by the point of organ placement.

### 2.1 Overexpression of Nodal and effects on laterality

The data above shows that when very early LR patterning steps are perturbed, and *Nodal* expression is incorrect, there can be a subsequent correction of this information downstream to lead to a lower number of tadpoles with reversed organs. However, we wanted to test whether direct misexpression of *Nodal* led to a clear mispatterning of laterality markers, and a reversal of organ *situs*, or whether misexpression of *Nodal* could also be corrected or tolerated by the developing embryo. What happens if *Nodal* is directly randomized, by forced misexpression of *Nodal* protein on the right side, or by overexpression of *Nodal* in general? A number of studies have already looked at the effect of Nodal overexpression in Xenopus, using both the injection of plasmid DNA [220, 221] and mRNA [222] (summarized in Table 10).

**Table 10:** Effect of xnr1 overexpression on Pitx2 laterality and organ situs.

| **Treatment** | **Incorrect**<br>***Pitx2* expression** | **Reversed Organs** | **Reference** |
| --- | --- | --- | --- |
| Xnr1 plasmids into 1 of 4 - left | **9%** |  | [220] |
| Xnr1 plasmids into 1 of 4 - right | **86%** |  | [220] |
| 50 pg Xnr1 into LPM dorsal blastomeres,<br>8-cell, bilateral | **85%** |  | [222] |
| DR injections xnr1 20pg pXEX |  | **61%** | [221] |
| DR injections xnr1 100pg pXEX |  | **71%** | [221] |
| DR injections xnr1 20pg pCSKAX |  | **76%** | [221] |
| DR injections xnr1 100pg pCSKA |  | **68%** | [221] |
| DL injections xnr1 20pg pXEX |  | **20%** | [221] |
| DL injections xnr1 100pg pXEX |  | **25%** | [221] |
| DL injections xnr1 20pg pCSKA |  | **33%** | [221] |
| DL injections xnr1 100pgpCSKA |  | **50%** | [221] |

However a study of the effects on all markers of laterality within the same group of treated embryos has not been carried out. Therefore we injected mRNA encoding *Xnr1*, the *Xenopus Nodal*, into one of two cells and studied the effect on the laterality of *Lefty* and *Pitx2* expression and scored the remaining tadpoles for organ *situs* (Table 11).

**Table 11:** Effects of Nodal overexpression on Lefty and Pitx2 laterality and organ situs and the degree of repair of incorrect laterality, calculated as the percentage of embryos with incorrect Nodal expression that have correct organ situs within the same clutches. R: right; Bi: bilateral; N: none. Positive degree of repair indicates that defects in early laterality markers are being corrected by the point of organ placement.

| **Incorrect**<br>***Lefty* expression** | **Incorrect**<br>***Pitx2* expression** | **Reversed Organs** | **Degree of Repair** |
| --- | --- | --- | --- |
| **61%**<br>(R: 27%, Bi: 32%, N: 2%) | **61%**<br>(R: 20%, Bi: 40%, N: 1%) | **39%** | **36%** |

While the misexpression of *Lefty* and *Pitx2* are consistent, the percentage of reversed organs drops dramatically. Embryos were coinjected with *β*-galactosidase mRNA and stained to compare the side of injection at the 2-cell stage with the phenotypes of *situs solitus* or heterotaxia (Table 12).

**Table 12:** Side of β-galactosidase expression compared with organ placement

| **Organ placement** | **Left-sided** | **Right-sided** | **Both sides** |
| --- | --- | --- | --- |
| ***Situs solitus*** | **158** | **70** | **28** |
| **Heterotaxia** | **22** | **100** | **34** |

There is a clear bias to right-sided *Nodal*-expression leading to misplacement of organs, but as Table 12 shows, the side of injection is not an absolute determining factor of the effect of *Nodal* overexpression on organ *situs*. Therefore it is possible that perturbation of *Nodal* levels can be fixed, as the side of injection and the level of marker misplacement versus the level of organ misplacement both show an incomplete dependence on the expression of *Nodal* to drive misplacement of organs.

## 3 Discussion

Gene-regulatory cascades form an important part of the middle phases of LR patterning [102], but important elements of physics and physiology lie upstream, in determining the sidedness of the first asymmetric transcription, and downstream, in implementing laterality cues toward asymmetric morpogenesis. Early events feed into different parts of the LR cascade, such as overexpression of Wnt8 and of the C314D mutant of mahogunin which is unable to ubiquitylate-tubulin. Indeed, some aspects of tissue morphogenesis may be intrinsic, using cell-level chirality to implement asymmetric looping directly, as may occur in *Drosophila* [96] and zebrafish [196].

While many organisms establish large-scale asymmetries, it is interesting that chirality is fundamental to individual cells an ancient evolutionary feature that metazoans exploit for macroscopic anatomical purposes. Aside from highlighting the propagation of properties across orders of magnitude of scale, laterality sheds light on the relationship between genome and fundamentally epigenetic factors. In single-cell ciliates, Beisson and Sonneborn demonstrated that reversal of a ciliary row in the cell cortex is propagated to offspring. Their function is also reversed, and cells eventually starve because food particles are swept into the wrong direction; their normal genome is powerless to rescue them [223, 224, 225, 226].

The original picture of the LR cascade made use of a midline barrier which separated distinct Left-and Right-sided programs of repression and induction [227]. This was subsequently revised by the finding that the L and R sides needed to communicate long-range via gap junction-mediated physiological signals for proper expression of early asymmetric genes [186, 228], but the importance of establishing a robust midline was clear [229, 230, 231, 232, 233]; indeed, the LR axis cannot be oriented without a midline that sets the axis of symmetry. While some animals are thought to establish the midline later than others, and the LR axis is often viewed as being defined after and with respect to the AP and DV axes, cleavage patterns and the prevalence of strictly bilateral gynandropmorphs resulting from very early cleavage events reveal that from insects to man [234, 235, 236], separating the L and R sides is one of the first things most embryos do [237]. Once the midline separates L and R compartments and they establish their unique identities, the standard model holds that cascades of genes become expressed sequentially, in a functional pathway that should propagate errors as genes inappropriately expressed on one side exert their effects and turn on/off downstream genes counter to their normal restricted unilateral patterns.

Most importantly, we have found, in both our new data and data from published experiments in *Xenopus*, discrepancies between the incidence of incorrect expression of early laterality markers and that of abnormal positioning of organs; these examples reveal the ability of embryos to correct defects in LR patterning over time (Figure 2). For example, early misexpression of unconventional myosin proteins (Table 2) and the mahogunin protein which targets α-tubulin for degradation (Tables 8,9) strongly disrupt the normal laterality of *Nodal* expression but have no effect on the positioning of organs in those embryos.

**Figure 2:**
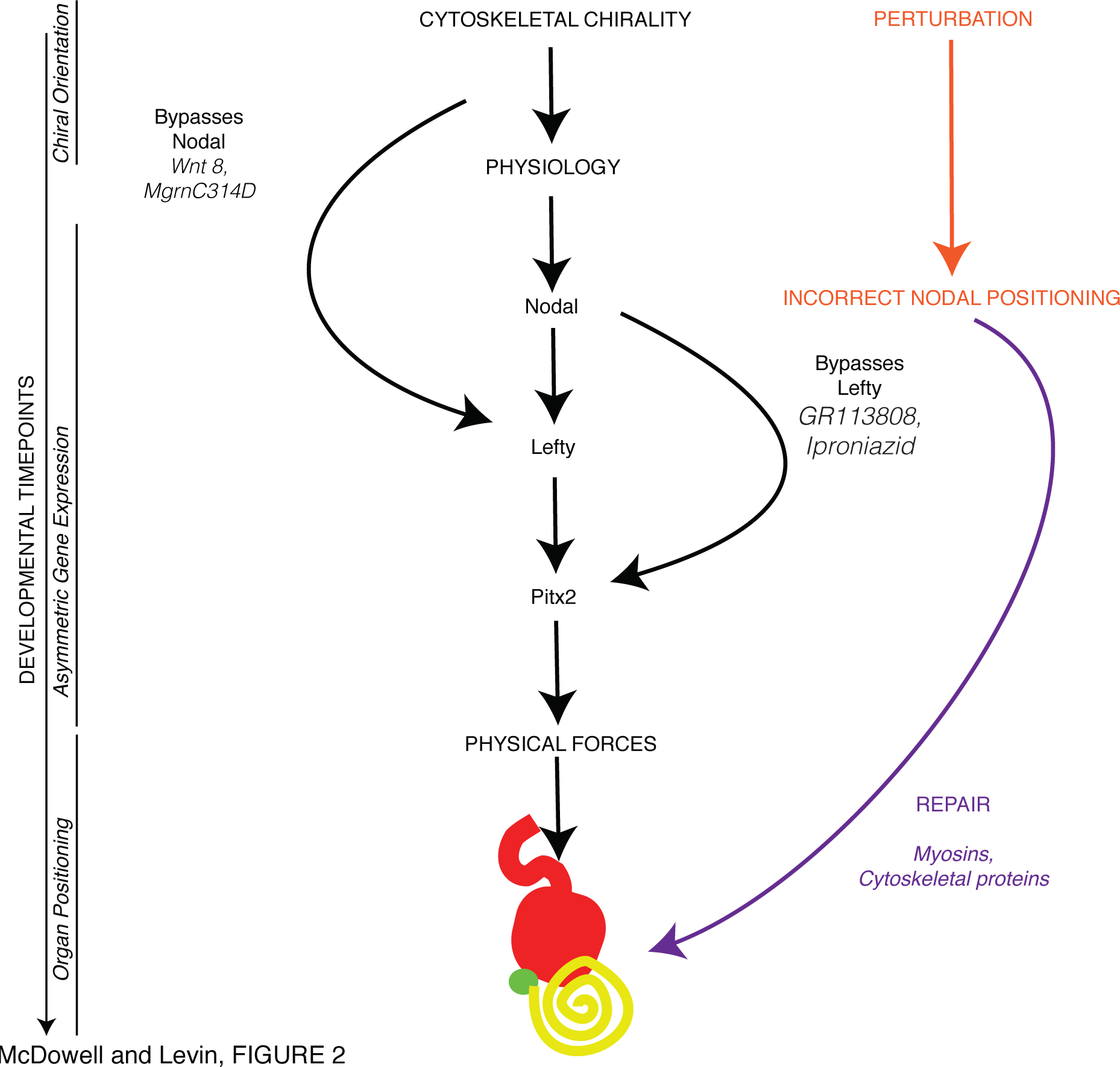
Summary of the LR pathway with possible points of fixing. The traditional linear pathway of LR patterning, from cytoskeletal chirality, physiological amplification through serotonin asymmetry and ciliary flow, expression of the laterality markers *Nodal, Lefty* and *Pitx2*, followed ultimately by physical bending to generate asymmetrical organ patterning is shown. Examples of data in the literature from experiments manipulating Wnt8 show that after the introduction of very early perturbations, early markers such as *Nodal* can appear normal, but later markers such as *Lefty, Pitx2* and organ *situs* can have incorrect laterality. This is also the case for the mahogunin mutant Mgrn C314D, whereas Mgrn wt overexpression has the opposite effect and results in incorrect *Nodal* expression, but correct *Lefty, Pitx2* and organ laterality. Likewise, Lefty laterality can be normal in cases where Nodal, Pitx2 and organ *situs* are incorrectly positioned with the serotonin receptor blocker Gr113808 and the monoamine oxidase inhibitor iproniazid. The identification of mechanisms that somehow detect abnormal sidedness of gene expression and institute corrections is one of the most exciting new vistas of the LR asymmetry field.

Our findings further cement the role of cytoskeleton as a conserved element of intracellular LR patterning: myosins, tubulins, and proteins regulating tubulin stability and actomyosin nucleation among species such as *Arabidopsis, Mollusca, Drosophila, Xenopus* and mouse are all functioning in asymmetry determination. However, the data reveal a much different picture than simply further details on cytoskeletal orienting machinery upstream of a linear asymmetric gene pathway. Induced errors in early asymmetry steps are sometimes fixed, or normalized, by the time of expression of subsequent downstream markers. This occurs most readily when very early steps are interfered with, but not for example when *Nodal* is directly misexpressed. However, in the case of *Nodal* misexpression, while the laterality of *Lefty* and *Pitx2* expression are incorrect, there is a reduction in the corresponding number of tadpoles with reversed organs (Table 11). One possibility is that the embryo needs time to notice and correct problems, or perhaps the early mechanisms are especially primed for correction.

In the case of low-frequency vibrations that induce LR patterning defects in early embryos, it appears that the early (cleavage) stages are crucial for enabling downstream repair mechanisms post-*Nodal*. Vandenberg *et al*. found that the level of *Nodal* expression randomization was constant and high despite the stage at which treatment began (as long as treatment occurred prior to blastula stage), but in all cases the percentage of resulting tadpoles with mispatterned organs was lower ([238], summarized in Figure 3A). This underscores the potential divergence between organ endpoints and marker expression in experiments with different timing conditions, and suggests that future studies cannot simply use gene expression readouts but must score organs as well to get a true picture of the effects of specific perturbations on the LR pathway. These data suggest that the earliest events (from 1-cell to st. 6) are important to enabling whatever mechanisms allow organs to form correctly despite abnormal *Nodal* laterality, as the ability to repair organ positioning increases progressively with treatments that start at later cleavage stages.

**Figure 3:**
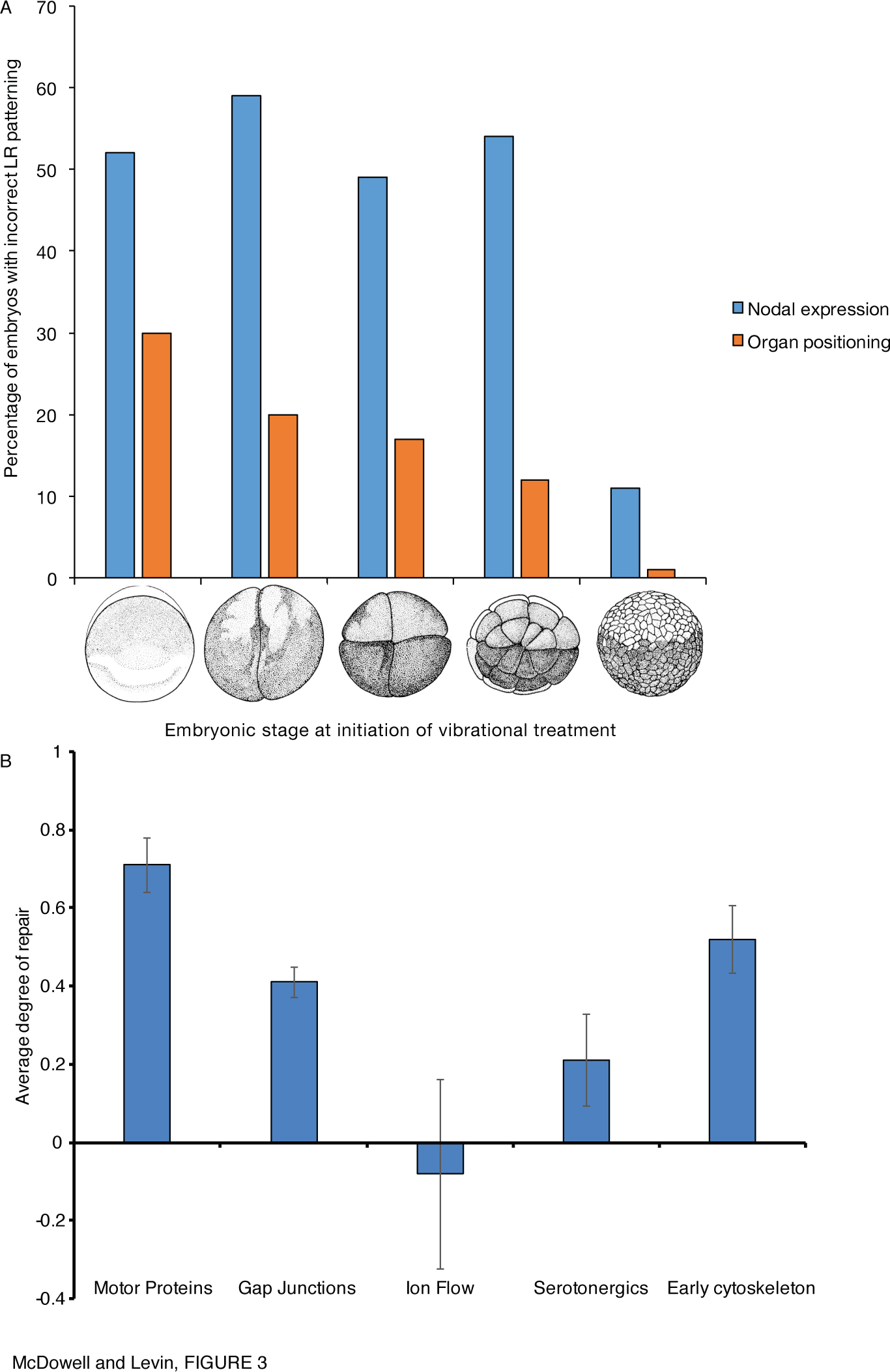
Variation in the ability to fix LR patterning. (A) Repair of early LR patterning defects induced by vibrational disrupton from early timepoints. Treatment of *Xenopus* embryos from the stages indicated to neurula (NF Stage 19) with low-frequency vibrations by Vandenberg *et al.* [238] caused LR patterning defects. Defects were observed both in the laterality of *Nodal* expression and in the positioning of organs, when pre-blastula mechanisms were targeted (but not later). However despite persistent vibrational disruption from the very earliest stages of embryo, defects in *Nodal* laterality were fixed by the point of organ positioning. Note the progressive increase of organ repair ability (as observed in reduction of organ defects despite high *Nodal* randomization) with treatments that start at each sequential cleavage. Images of *Xenopus* embryos Copyright 1994 Pieter D. Nieuwkoop and J. Faber [239]. (B) The ability of embryos to rescue LR patterning defects depends on the target of experimental perturbation. Experiments analyzed throughout this study were grouped according to their functional targets and the average degree of repair (difference between incorrect organ *situs* and incorrect *Nodal* expression sidedness) was plotted for each group (see Supplementary Table 1). Error bars shown are the standard error of the mean among different treatments as reported in the studies cited above.

Errors in asymmetrically-expressed genes that were induced by perturbing myosin, tubulin and gap junctional communication are subsequently repaired, while those in Wnt expression and ion-channel regulation are not. It is currently unknown why this is the case. An obvious possibility is the existence of parallel pathways a sort of parity check, that allows the embryo to test whether the sidedness of specific gene products is correct or not. The details remain to be characterized. It should be noted however that another potential layer of complexity in these stochastic data could be due to the possibility that different embryos from the same batch might be relying on somewhat different mechanisms for LR asymmetry (discussed in detail in [240]).

Interestingly, grouping the experimental results on normalization of downstream steps by the type of early perturbation (Figure 3B, Supplementary Table 1) suggests that the degree of laterality repair capability varies depending on the type of perturbation (the LR pathway target that was perturbed to generate the *Nodal* randomization). Perturbations of motor proteins, and the early cytoskeleton, within the first cell division, is tolerable to a certain degree as it largely becomes normalized by the time of organogenesis; but manipulations of ion flows are apparently not possible to recover from. It should also be noted that in most cases within the serotonergic group, *Lefty* appeared to be bypassed (Table 7). These data suggest that further study into the different degree of pathway repair downstream of targeting different types of components of LR patterning may shed greater light on the mechanism, and robustness, of LR patterning and the normalization of downstream targets despite randomized prior steps.

The data also reveal non-linearity of the pathway as some elements can be bypassed: interfering with early serotonin and monoamine oxidase signaling results in organ-level LR defects despite the correct expression of *Lefty* (Table 7). A dynamical systems view of these repair pathways, as a network rather than a linear pathway with necessary and sufficient master regulators, is necessary, as has already been noticed in the field of cancer and the search for driver genes [241, 242, 243, 244, 245, 246]. The future is likely to involve not only molecular-genetic picture of these repair pathways, but a cybernetic, systems-control view of the information processed by these closed-loop, shape-homeostatic capabilities of embryogenesis [247, 248, 249, 250, 251]. It thus becomes clear that checking immediate downstream consequences of gene misexpression is not sufficient for functional analyses; a subtler strategy targeting further upstream of a gene of interest, which gives the embryo more time to recognize errors, is required to form a full picture of regulatory networks and necessary and sufficient claims for an explanation of what gene product determines expression of some other gene product.

In a sense, pattern homeostasis, such as seen in reparative regeneration widely across the tree of life [210], could be seen as a primary biological capability. Development is really just an example of regeneration restoring the whole body from 1 cell (fertilized egg). No wonder that regenerative repair, which can make up for disruption of pattern with flexible corrective processes, also occurs in embryogenesis. While not surprising given regulative development as a whole, it sheds new light on the definition of master regulator or determinant genes for specific developmental outcomes. While loss-and gain-of-function tests (such as those that had been performed for *Nodal* in the early years of the study of the LR pathway) may indicate that a certain genes expression is a driver of what happens next, it has to be kept in mind that this may not be the whole story. Two things can be readily missed by experiments that are narrowly focused on direct functional change of the gene expression and an immediate readout of direct downstream response genes. First, the downstream consequences could be wiped out by corrective mechanisms, which can limit the validity of necessary and sufficient claims for specific gene products with respect to final patterning outcome. Second, the results may be quite different if that genes expression is deranged by targeting much earlier steps (giving the organism time to notice the problem and activate robust repair pathways).

These data and analyses suggest a new direction in this field, focused on a systems-level understanding of how distinct molecular steps are multiplexed by the embryo for monitoring, setting, and continuously re-setting the LR identity of its tissues. In the sense that embryogenesis is an example of the more general phenomenon of regeneration (rebuilding the whole body from 1 cell), the LR asymmetry mechanisms may be a window on a much more general and widely-relevant phenomenon than just the patterning of the LR axis. The study of these dynamics could reveal fundamental aspects of how genetics interfaces with physics to implement robust, self-correcting systems. The impact of this would extend beyond development and birth defects, to the understanding of evolution, regenerative medicine, and artificial life.

## 4 Materials and Methods

### 4.1 Cloning

Subcloning was carried out into pCS2+ using standard methods as described previously [112].

### 4.2 Animal Husbandry

This study was carried out in strict accordance with the recommendations in the Guide for the Care and Use of Laboratory Animals of the National Institutes of Health and Tufts IACUC protocol #M2014-79. *Xenopus* embryos were collected and maintained according to standard protocols [252] in 0.1 x Modified Marcs Ringers (MMR), pH 7.8, and staged according to [239].

### 4.3 Microinjection

Capped, synthetic mRNAs were dissolved in water and injected into embryos in 3% Ficoll using standard methods [252]. mRNA injections were made into the animal pole of eggs within 30 minutes post fertilization (mpf) at 14 °C, or into 1 of 2 cells of Stage 2 [239] embryos (as indicated) using borosilicate glass needles calibrated to deliver a 10 nl injection volume.

### 4.4 Laterality Assays

*Xenopus* embryos were analyzed for position (*situs*) of three organs; the heart, stomach and gallbladder [186] at Stage 45 [239]. Heterotaxic embryos were defined as having a reversal in one or more organs. Only embryos with normal dorsoanterior development and clear left-or right-sided organs were scored. A χ^2^ test was used to compare absolute counts of heterotaxic embryos.

### 4.5 *In situ* hybridization

Whole mount *in situ* hybridization was optimized using standard protocols [253, 254] with probes against *Xnr1* (the *Xenopus Nodal*) [221], *Lefty* [255] and *Pitx2* [222] generated *in vitro* from linearized template using DIG labeling mix (Roche). A χ^2^ test was used to compare absolute counts of embryos with correct (expression on the left lateral plate mesoderm) versus incorrect (absent, bilateral or right-sided) marker expression.

## 5 Acknowledgements

We thank Joan Lemire and Jean-Francois Par for cloning assistance and helpful comments on the manuscript, and other members of the Levin lab and the LR asymmetry and regeneration communities for many helpful discussions. We gratefully acknowledge support of the Templeton World Charity Foundation TWCF0089/AB55, sub award from Physical Science Oncology Center supported by Award Number U54CA143876 from the National Cancer Institute, and the Paul G. Allen Family Foundation.

## 6 Comments

Comments can be made by contacting the corresponding author at or on the version of this preprint archived at *The Self-Journal of Science*:http://sjscience.org/article?id=536

## 7 Supplementary figures

- Supplementary Table 1: Experimental evidence for fixing grouped according to functional targets.

